# Games and the treatment convexity of cancer

**DOI:** 10.1101/2023.02.27.530257

**Authors:** Péter Bayer, Jeffrey West

**Affiliations:** Toulouse School of Economics and Institute for Advance Study in Toulouse, 1 Esplanade de l’Université, 31080 Toulouse, France; Department of Integrated Mathematical Oncology, Moffitt Cancer Center, 12902 USF Magnolia Drive, Tampa, FL 33612, United States

**Keywords:** Game theory, cancer, dilemma strength, treatment convexity

## Abstract

Evolutionary game theory has been highly valuable in studying frequency-dependent selection and growth between competing cancer phenotypes. We study the connection between the type of competition as defined by properties of the game, and the convexity of the treatment response function. Convexity is predictive of differences in the tumor’s response to treatments with identical cumulative doses delivered with different variances. We rely on a classification of 2 × 2 games based on the signs of ‘dilemma strengths’, containing information about the kind of selection through the game’s equilibrium structure. With the disease starting in one game class, we map the type of effects treatment may have on the game depending on dosage and the implications of treatment convexity. Treatment response is a linear function of dose if the game is a prisoner’s dilemma, coordination, or harmony game and does not change game class, but may be convex or concave for anti-coordination games. If the game changes class, there is a rich variety in response types including convex-concave and concave-convex responses for transitions involving anti-coordination games, response discontinuity in case of a transition out of coordination games, and hysteresis in case of a transition through coordination games.

## 1 Introduction

Game theory is the mathematical discipline and universal language of strategic interaction. Pioneered by John von Neumann and Oskar Morgenstern, then further developed by John Nash, the field was originally created with the goal of formalizing and understanding the behavior of rational decision-makers, making it a mainstream tool of modern economics.

In the 1960s and 1970s, theoretical biologists William D. Hamilton, George R. Price, and John Maynard Smith were creating a formal model of evolution by natural selection. Recognizing the potential offered by game theory in modeling frequency-dependent selection within biological populations, they adopted elements of non-cooperative game theory which, combined with novel tools of game dynamics, formed the core of the newly developed field of evolutionary game theory. Hamilton’s rule of cooperation arising from kin selection [1] is stated in terms of a two-player prisoner’s dilemma game, while Maynard Smith and Price’s theory of animal conflict [2] makes explicit use of game theory. The latter work, based on Hamilton’s ‘unbeatable strategy’, develops the concept of the ‘evolutionary stable strategy’ (ESS) and is seen as the starting point of evolutionary game theory.

The emergence and progression of cancer is driven by Darwinian evolution by natural selection, making evolutionary game theory one of the key tools used by the novel field of mathematical oncology [3]. Starting with [4], interactions between cancer cells were modeled using evolutionary game theory, yielding important insights into the selection pressures of cell phenotypes and the frequency-dependent growth of the cancer population (for an overview, see [5]). The theory’s insights helped challenge long-standing paradigms of cancer treatment [6], inspire novel approaches to cancer therapy [7, 8], which then lend themselves to be adjusted and applied in specific cancers [9], and, finally, clinical trials [10], making their way towards medical practice.

In this paper we propose a link between frequency-dependent competition of cancer cell phenotypes as described by an evolutionary game and the ‘treatment convexity’ of cancer, that is, the differences in response to treatment schedules with identical cumulative dose levels but different dose variances. While these ‘second-order effects’ have been shown to be null in theoretical models with a single cancer phenotype [11], as we will show, framing the selection and growth of cancer as a competition of as few as two cancer cell phenotypes engaged in an evolutionary game, a rich variety of nonlinear second-order effects are obtained. In this paper we show these effects to be dependent upon the type of game the phenotypes are engaged in and how applying therapy in different doses changes that game class.

### 1.1 Game classification and dilemma strength

Various taxonomies exist for non-cooperative 2×2 games following different criteria. A classic one, introduced for symmetric non-cooperative games with no Pareto-dominant strategy profile and no strongly stable equilibria (leaving only four games) is through the directional and comparative payoff effects imposed on self and partner by a unilateral switch of strategy from the game’s minimax outcome [12].^1^. A complete classification, [13] creates a periodic table of all 144 strictly ordinal 2 × 2 non-cooperative games by their topological and algebraic properties (a taxonomoy of non-strict ordinal games has been created by [14]).

Static evolutionary games are mathematically equivalent to symmetric normal-form non-cooperative games, hence, their classification is simpler. Being characterized by four numbers, there are a total of 4! = 24 strict ordinal 2 × 2 symmetric games.^2^ Accounting for equivalence up to a relabeling of both players’ strategies, we get a total of 24/2 = 12 games.

In this paper we use a classification based on [15], leveraging two parameters known as ‘dilemma strengths’ [16]. These parameters obtain as the payoff differences of a player between her two possible strategies while keeping the opponent’s strategy fixed (and, possibly, normalized as in [17]). The signs of these parameters define four quadrants in 2-dimensional space, giving four game classes.

In social dilemmas with strategies labeled as ‘cooperate’ and ‘defect’, the two parameters represent the strength of the so-called ‘gamble-intending-dilemma’ (the payoff gain of defecting against a cooperating opponent instead of cooperating) and ‘risk-averting-dilemma’ (the payoff gain of defecting against a defecting opponent instead of cooperating). As our strategies will be labeled 1 and 2 with no social dilemma interpretation attached to them, we find it convenient to use parameters Δ_1_, denoting the payoff advantage of playing 1 instead of 2 against an opponent with strategy 1, and Δ_2_, the payoff advantage of playing 2 instead of 1 against an opponent with strategy 2. We nonetheless keep the labels of the four classes defined by the signs of these two parameters, as used in the social dilemma strength literature (with the labels themselves originally being the names of well-known classes of 2 × 2 games).^3^ The signs of these two parameters provide all information about the game’s best-response and equilibrium structure. If both Δ_1_ < 0 and Δ_2_ > 0, strategy 2 (“defect”) is dominant and the game is a prisoner’s dilemma. If Δ_1_ > 0 and Δ_2_ < 0, strategy 1 (“cooperate”) is dominant and the game is harmony (so-called for an absence of a social dilemma, a.k.a. “deadlock” and “anti-Prisoner’s dilemmma”). If both are positive, both strategies obtain as stable equilibria, and the game is a coordination game (a.k.a. “stag hunt”). If both are negative, the only stable equilibrium is mixed and the game is an anti-coordination game (a.k.a. “hawk-dove”, “chicken”, and “snowdrift”).

### 1.2 Measuring effective games in oncology

Evolutionary game theoretic models have played an important role in characterizing the range of possible cell-cell interactions at play in cancer progression and treatment [18, 19, 5]. Exciting progress has been made recently in quantifying game-theoretic interactions between cell types in cancer. For example, public goods games have been observed in engineered defector cells [20] and for IGF-II growth factor production in neuroendocrine pancreatic cancer [21]. Using Lotka-Volterra competition models, others have shown experimentally the influence of competitive interactions between sensitive and resistant cell lines on the optimal between metronomic schedule (low doses of drug delivered at high frequencies) can control tumor cell dynamics better than a high dose schedule [22]. Similar models quantified the resource-dependent (low/high pH and low/high glucose) competition dynamics between two breast cancer cell lines [23]. Another predator-prey mathematical model showed evidence of reciprocal antagonism at play within competition dynamics of parental and radiation-resistant cell lines [24]. These models have also been applied to endocrine therapy [25] and doxorubicin therapy [26, 27] in breast cancer cell lines. There is also evidence of the existence of positive or cooperative ecological interactions in cancer: one recent study indicated paracrine signaling as a mechanism for which the dominant cell line’s growth is enhanced by the presence of a co-cultured second cell line [28]. Evolutionary game theory models have also been used to quantify competition dynamics between multiple myeloma and bone marrow cells within an experimental microecology setup, with a stable drug gradient between connected microhabitats [29].

Recent literature has also shown that cancer treatment can induce a transition of the game class describing the underlying dynamics. For example, co-cultured non-small cell lung cancer parental and gefitinib-resistant cell lines transitioned from a harmony game in untreated culture (parental dominates) to a prisoner’s dilemma game in etoposide treated conditions (resistant dominates) [30]. Similarly, co-cultured parental and alectinib-resistant cell lines transition from an anti-coordination game (when cultured with cancer-associated fibroblasts) to a harmony game under alectinib treatment [31].^4^

Surveying past literature clearly shows that a wide range of game class dynamics are relevant in cancer. Previous work on the game theory in cancer often focuses on the relevancy of specific classes of games (e.g. prisoner’s dilemma [32, 33] or coordination games [34]), linking properties of the game class in question to properties of the disease. Others focused on mapping the parameter space of games of general form through exhaustive simulation with a lesser focus on the games’ properties [35].

### 1.3 Second-order treatment effects

As treatment is administered to the patient, the eco-evolutionary dynamics of cancer changes. The treatment’s primary effect is the reduction of cancer cells’ reproductive potential, expressed by the fitness game’s payoff values, leading to a reduction of cancer’s proliferative potential upon receiving the treatment. The secondary effect is the changed selection dynamics between competing cancer phenotypes, expressed by the changes of the dilemma strength parameters. If the parameters change sign as a result of intervention, there is a qualitative change in the tumor’s selection dynamics.

Even with precise characterization of the selection dynamics at play, it is not a straightforward task to design an optimal treatment schedule. We propose a framework based on the principles of convexity, and apply these principles to game theoretic models. A given treatment protocol has an associated mean dose (first-order metric) and dose variance (second-order metric). In clinical practice, much attention is given to maximizing first-order treatment effects by decreasing the amount of time over which a cumulative dose is delivered, as toxicity allows [36].

Here, we restrict our analysis to treatment schedules with identical cumulative dose and illustrate the benefits achieved by altering only the dose variance. Examples are shown in Figure 1A. Consider an “even” dosing schedule consisting of a constant, periodic dose delivered: 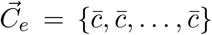 (see Figure 2A, purple). We compare this to an “uneven” dosing schedule with identical cumulative dose that alternates between a high and low dose: 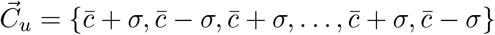.

**Figure 1.**
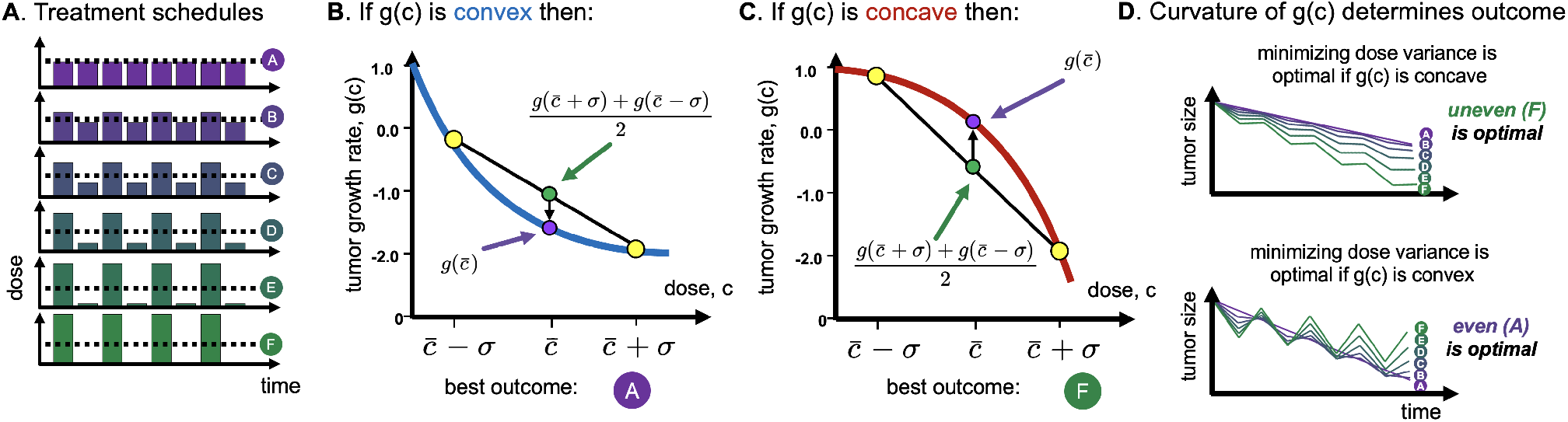
Convexity leads to differential outcomes. (A) example treatment schedules with identical cumulative dose, ranging from “even” (purple) to “uneven” high/low dosing. (B) By Jensen’s inquality, if tumor growth rate, *g*(*c*), is convex, then even dosing will minimize growth rate. (C) Similarly, if *g*(*c*) is concave, uneven dosing will minimize growth rate. (D) Example trajectories under concave (top) or convex (bottom) functions.

**Figure 2.**
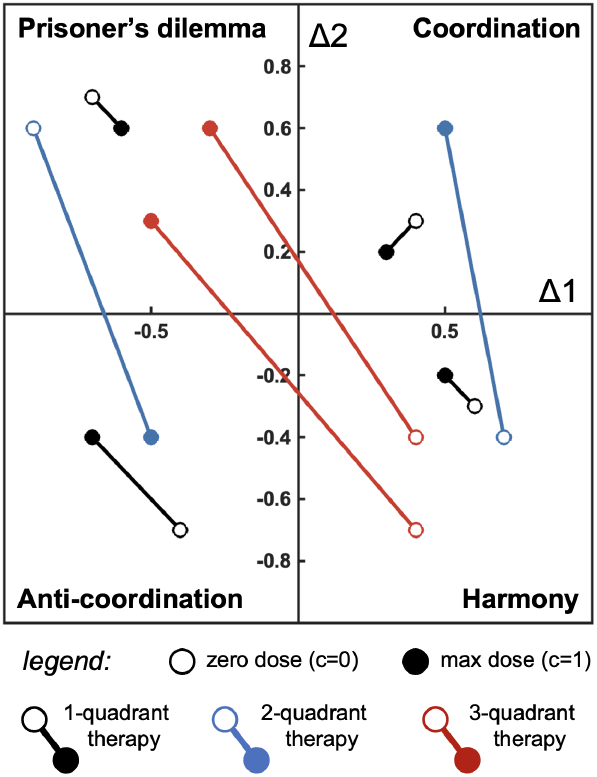
Some possible game class transitions.

We compare this suite of treatment schedules to determine which will minimize the tumor growth rate. It has previously been shown that the tumor-minimizing optimal schedule (even versus uneven) depends on the curvature of growth dynamics [37, 38]. By Jensen’s inequality, the negative growth rate for even schedule, 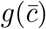, is lower than the average growth rate of the high and low dose uneven strategy, 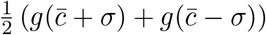 (compare purple and green dot in Figure 1B). The converse is seen in Figure 1C, illustrating that uneven schedules minimize tumor growth for convex functions. Example tumor trajectories are shown in Figure 1D.

Convexity has been shown to be predictive in vitro [38]. However, previous work does not consider the influence of cell-cell interactions. We extend convexity analysis to a system of two interacting populations competing according to a fitness landscape described by an evolutionary game. The model framework introduced in the next section displays a remarkable range of dynamical behavior including linearity, convexity or concavity (local and/or global), and hysteresis. Broadly, these phenomena are determined by game class.

## 2 Methods

### 2.1 A 2 × 2 game driving selection

As a minimum requirement for game theoretic analysis, we consider a tumor consisting of two strategies (e.g. phenotypes or cell lines) of cancer cells, 1 and 2. We assume that the two types interact within the tumor microenvironment and that the cancer cells’ reproduction is dependent upon these interactions. We assume that interactions are pairwise. The strict game-theoretic interpretation is that cancer cells exist in a well-mixed population (tumor), are matched into pairs randomly and interact according to their types, then realize payoffs (reproduction) as per a normal-form game which we denote by **Π**. A type *i* cell interacting with a type *j* cell receives payoffs *π*_*ij*_ ≥ 0 (Table 1). Without loss of generality we assume *π*_11_ ≥ *π*_22_, that is, we label the types such that type 1 grows fastest in its own environment; we therefore call type 1 the growth-optimal strategy of cancer.

**Table 1.** Left: the growth game, **Π**, governing the growth and the interaction of cancer cell types absent therapy. Right: the treatment game, **Γ** = **Π** − *c***K**.

|  |  | Environment |  |  |  | Environment |  |
| --- | --- | --- | --- | --- | --- | --- | --- |
|  |  | Type 1 | Type 2 |  |  | Type 1 | Type 2 |
| Focal cell | 1 | $\pi_{11}$ | $\pi_{12}$ | Focal cell | 1 | $\pi_{11} - c\kappa_{11}$ | $\pi_{12} - c\kappa_{12}$ |
| | 2 | $\pi_{21}$ | $\pi_{22}$ | | 2 | $\pi_{21} - c\kappa_{21}$ | $\pi_{22} - c\kappa_{22}$ |

Therapy is modeled through its effects on the payoff parameters on the growth game **Π**. To capture the pure effect of frequency dependence on treatment convexity, we assume that treatment reduces the growth parameters proportionally to the the dosage of therapy. For *i, j* ∈ {1, 2}, let *κ*_*ij*_ ≥ 0 be the maximum reduction of growth parameter *π*_*ij*_. The game under treatment dose *c* is denoted by **Γ**(**Π, K**, *c*), shortened to **Γ**(*c*) for ease of notation whenever convenient, with its payoff components given by *γ*_*ij*_(*c*) = *π*_*ij*_ − *cκ*_*ij*_.

### 2.2 Game classes

Define

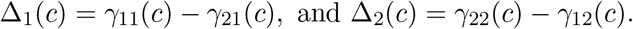

For type *i* ∈ {1, 2}, Δ_*i*_(*c*) is the evolutionary advantage of type *i* against the opposite type in type *i*’s environment, given dosage *c*. We distinguish four types of games based on the signs of these two parameters (Table 2).

**Table 2.**
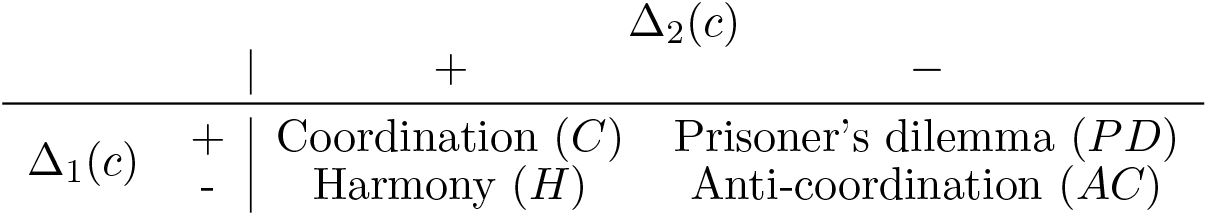
The classification of the interaction game **Γ**(*c*) depending on the signs of Δ_*i*_(*c*).

| | | $\Delta_2(c)$ | |
| --- | --- | --- | --- |
|  |  | + | - |
| $\Delta_1(c)$ | + | Coordination ($C$) | Prisoner's dilemma ($PD$) |
| | - | Harmony ($H$) | Anti-coordination ($AC$) |

In a prisoner’s dilemma, the growth-optimal strategy is dominated, coordination games are characterized by bi-stability of the two strategies, in anti-coordination games, evolution selects a mix of the two cell types, while in a harmony game, the growth-optimal strategy is dominant. As the growth game **Π** represents the status quo without intervention from the treating physician, we label the four quadrants based on that game as *PD, C, AC*, and *H*.^5^

### 2.3 Equilibrium growth rates

Let *G*: ℝ^2×2^ × ℝ^2×2^ × ℝ → 𝒫 (ℝ) be the correspondence mapping games **Γ**(**Π, K**, *c*) to their stable equilibrium payoffs. Then,

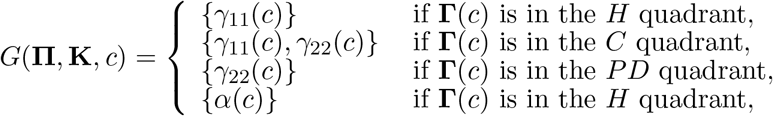

where

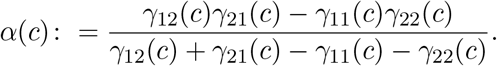

The derivation of *G* in the *H, C*, and *PD* quadrants is straightforward: In the *H* quadrant, the only equilibrium strategy is pure type 1, in the *PD* it is the pure type 2. In the *C* quadrant, both pure type 1 and type 2 are stable equilibria with an unstable mixed equilibrium characterized by

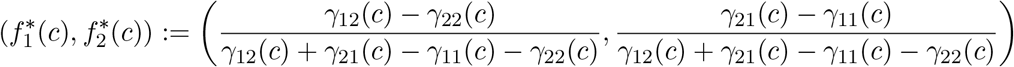

separating the basins of attraction between them. In the *AC* quadrant, the unique equilibrium frequency vector is given by 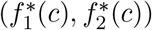, identical to the unstable, mixed equilibrium in the *C* quadrant. The equilibrium payoff is the sum of equilibrium frequencies weighted by payoff values: 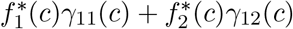 (or equivalently, 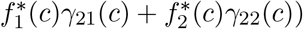, which simplifies to *α*(*c*).

As with **Γ**, we shorten the notation to *G*(*c*) whenever convenient and does not cause confusion. Treatment convexity will be determined by the convexity of (the component functions of) *G*(*c*).

### 2.4 Threshold doses leading to changes in game class

Of interest are growth and treatment parameters that result in a change of game class. Figure 2 shows possible game class transitions. The game representing treatment under maximum tolerable dose (solid circle) may lie within the same quadrant as the growth game under zero dose (open circle) as seen in black lines, or may transition to a neighboring quadrant (blue lines), or span three quadrants (red lines). To quantify the doses effecting quadrant transitions, we define *c*_1_ and *c*_2_ as

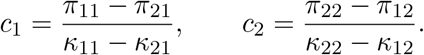

If *c*_1_, *c*_2_ ∈ [0, 1] then, they are the dose values for which Δ_1_(*c*_1_) = 0 and Δ_2_(*c*_2_) = 0, respectively. If both are outside the (0, 1) interval, then treatment spans only one game quadrant, if one is in (0, 1) it spans two, if both are in (0, 1), it spans three quadrants.

### 2.5 Population dynamics

While we base our analytics purely on the selection dynamic as defined by the game Γ(*c*), we conduct simulations in a dynamic model capturing both population and selection effects. For *i* = 1, 2 let *x*_*i*_(*t*) denote the population of cell type *i* at time *t* ≥ 0. Let *x*(*t*) = *x*_1_(*t*) + *x*_2_(*t*) denote the tumor’s size and *f*_*i*_(*t*) = *x*_*i*_(*t*)*/x*(*t*) denote the frequencies of the cell types. The growth rates of each phenotype is given as *π*_*i*_(*t*) =∑_*j*∈{1,2}_ *π*_*ij*_(*t*)*f*_*j*_(*t*). The model’s variables, parameters, and equations are shown in Table 3. The variables’ time dependence is omitted for simplicity whenever we think it does not cause confusion.

**Table 3.** The ingredients and equations of describing a model of growth governed by the game **Γ**(*c*).

| Variables |  | Parameters |  |
| --- | --- | --- | --- |
| $x_i(t)$ | Population sizes | $\pi_{ij}$ | Growth parameters |
| $x(t)$ | Tumor size ($x_1(t) + x_2(t)$) | $\kappa_{ij}$ | Treatment parameters |
| $f_i(t)$ | Frequencies ($x_i(t)/x(t)$) | $c(t)$ | Treatment dose |

## 3 Results

### 3.1 Convex and linear growth dynamics by game class

One classical mathematical formulation that is commonly-used to model cancer treatment response is the Norton-Simon model[11, 39]. This model assumes that the effect of treatment is linearly proportional to instantaeous growth rate: 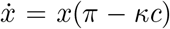 for *π, κ >* 0. It is a well-known result that tumor size is proportional to the cumulative dose delivered, 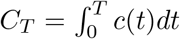 [40]. To re-phrase: second-order effects are null in this model formulation. Rearrangement of dosing to be front-loaded, back-loaded, or evenly spread all lead to identical outcomes. An argument for this result can be made solely from the linearity of the growth function: Because growth is a linear function of dose, all schedules with equivalent cumulative dose lead to equivalent outcomes.

Two example therapies are shown in Figure 3A: prisoner’s dilemma (black) and anti-coordination game (red). The equilibrium growth dynamics in Figure 3B of the prisoner’s dilemma are a linear function of dose, indicating by convexity theory [37] that neither even (Figure 2C, purple) nor uneven (Figure 2C, green) will outperform. This is confirmed in panel D, where all schedules perform approximately equally^6^. In contrast, the anti-coordination equilibrium growth dynamics are strongly convex, leading the continuous therapy (Figure 2E, purple) to outperform all other schedules.

**Figure 3.**
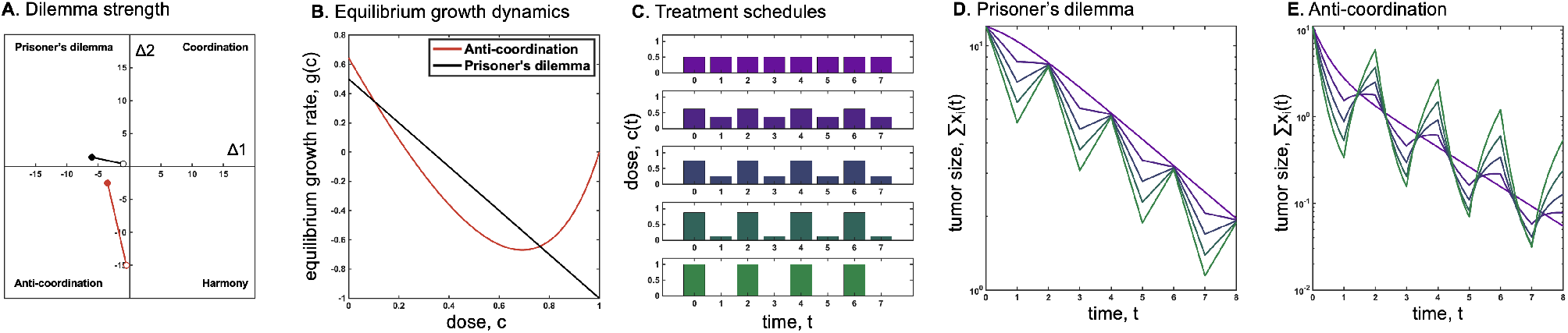
Convexity predicts differential outcomes. (A) Example treatment dynamics: prisoner’s dilemma (black) and anti-coordination (red). (B) Equilibrium growth dynamics, shown for Prisoner’s dilemma (black; **Π** = [1, 0.1, 2, 0.5]; **K** = [5, 2.5, 0, 1.5]) and Anti-coordination (red; **Π** = [0, 20, 0.5, 5]; **K** = [7, 15, 4, 2.5]). (C) Each game is subject to a suite of treatment schedules with identical cumulative dose, where 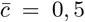; *σ* ∈ [0, 0.125, 0.25, 0.375, 0.5]. (D) Even treatment (purple) minimizes tumor volume for the anti-coordination game, matching the prediction based on convex growth dynamics in panel B. (E) All schedules lead to approximately equivalent dynamics for the prisoner’s dilemma game, matching the prediction based on linear growth dynamics in panel B.

In the next sections, we showcase ways through which different game classes and the transitions between them affect the convexity of treatment response with respect to treatment dose.

### 3.2 Convex and concave anti-coordination games and changing convexity from transitions

While equilibrium growth is a linear functions of dose in three of the four quadrants (PD, H, C), this is not the case for anti-coordination games. To obtain a sufficient and necessary condition for the convexity of *α*(*c*), rewrite:

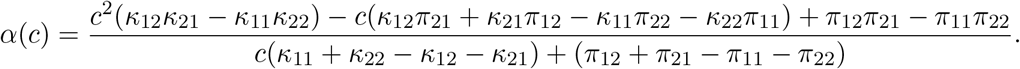

The second derivative is given by:

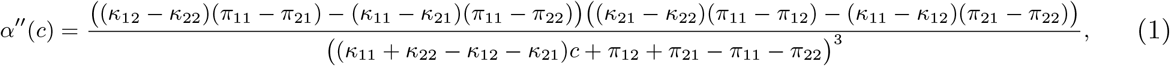

The numerator is independent of *c*, while the denominator is always positive as **Γ**(*c*) is an anti-coordination game, hence *α*(*c*) is convex if and only if the numerator in expression in (1) is positive. As it is independent of dosage, **Π** and **K** exactly determine convexity.

Upon transitioning into the *AC* quadrant from a neighboring quadrant *PD* or *H*, an additional component plays a role: the discontinuity of the slope of the equilibrium growth rate at the point of transition: if slope is higher pre-transition, equilibrium growth will be a convex function of dose around the transition point, while if it lower, the function will be convex. The combination of these two effects produces a total of four cases that is summarized by the following conclusion: if the growth-treatment game defines a transition into (or, similarly, out of) the *AC* quadrant, equilibrium growth may be a convex, concave, convex-concave, or a concave-convex function of treatment dose.

Stated formally, assume that **Π** lies in the *H* quadrant and **Π** − **K** in the *AC* quadrant. Then, Γ(*c*) is in *H* for *c < c*_1_ and *AC* for *c > c*_1_.

Equilibrium growth rate is well-defined, so *G*(*c*) is single valued. Let *g*(*c*): [0, 1] → ℝ be the equilibrium growth-rate function given by

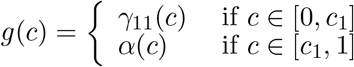

The function *g*(*c*) is linear and decreasing over the [0, *c*_1_] interval and, since *α*(*c*_1_) = *γ*_11_(*c*_1_), it is a continuous function over all of [0, 1].

Then, *g*(*c*) is

1. concave-convex if *α*′(*c*_1_) *< κ*_11_ and *α*′′(*c*) > 0,
2. convex if *α*′(*c*_1_) *> κ*_11_ and *α*′′(*c*) > 0,
3. concave if *α*′(*c*_2_) *< κ*_22_ and *α*′′(*c*) < 0,
4. convex-concave if *α*′(*c*_2_) *> κ*_22_ and *α*′′(*c*) < 0.

If **Π** is in the *PD* quadrant, the same analysis applies with *c*_2_, *γ*_22_(*c*), and *κ*_22_. Table 4 showcases three example sets of growth and treatment parameters that produce different convexity properties in transitioning into the *AC* quadrant, while Figure 4 showcases three possible tumor’s equilibrium growth functions in these games.

**Table 4.** Three sets of growth and treatment parameters defining transitions into the *AC* quadrant.

| $\mathbf{\Pi}_1^{\text{HA}}$ | 1 | 2 | $\mathbf{\Pi}_2^{\text{HA}}$ | 1 | 2 | $\mathbf{\Pi}^{\text{PA}}$ | 1 | 2 |
| --- | --- | --- | --- | --- | --- | --- | --- | --- |
| 1 | 7 | 35 | 1 | 8 | 1 | 1 | 12 | 0 |
| 2 | 4 | 6 | 2 | 5 | 0 | 2 | 14 | 4 |
| $\mathbf{K}_1^{\text{HA}}$ | 1 | 2 | $\mathbf{K}_2^{\text{HA}}$ | 1 | 2 | $\mathbf{K}^{\text{PA}}$ | 1 | 2 |
| 1 | 14 | 30 | 1 | 6 | 3 | 1 | 4 | 2 |
| 2 | 8 | 2 | 2 | 0 | 4 | 2 | 20 | 10 |

**Table 5.** Three sets of growth and treatment parameters. They all define transitions between the PD and H quadrants via the AC quadrant but the minimum of equilibrium growth obtains in different quadrants.

| $\Pi_1^{CH}$ | 1 | 2 | $\Pi_2^{CH}$ | 1 | 2 | $\Pi_3^{CH}$ | 1 | 2 |
| --- | --- | --- | --- | --- | --- | --- | --- | --- |
| 1 | 8 | 6 | 1 | 12 | 2 | 1 | 6 | 1 |
| 2 | 0 | 7 | 2 | 6 | 4 | 2 | 3 | 2 |

| $K_1^{CH}$ | 1 | 2 | $K_2^{CH}$ | 1 | 2 | $K_3^{CH}$ | 1 | 2 |
| --- | --- | --- | --- | --- | --- | --- | --- | --- |
| 1 | 10 | 3 | 1 | 9 | 3 | 1 | 10 | 3 |
| 2 | 11 | 5 | 2 | 11 | 5 | 2 | 11 | 5 |

**Figure 4.**
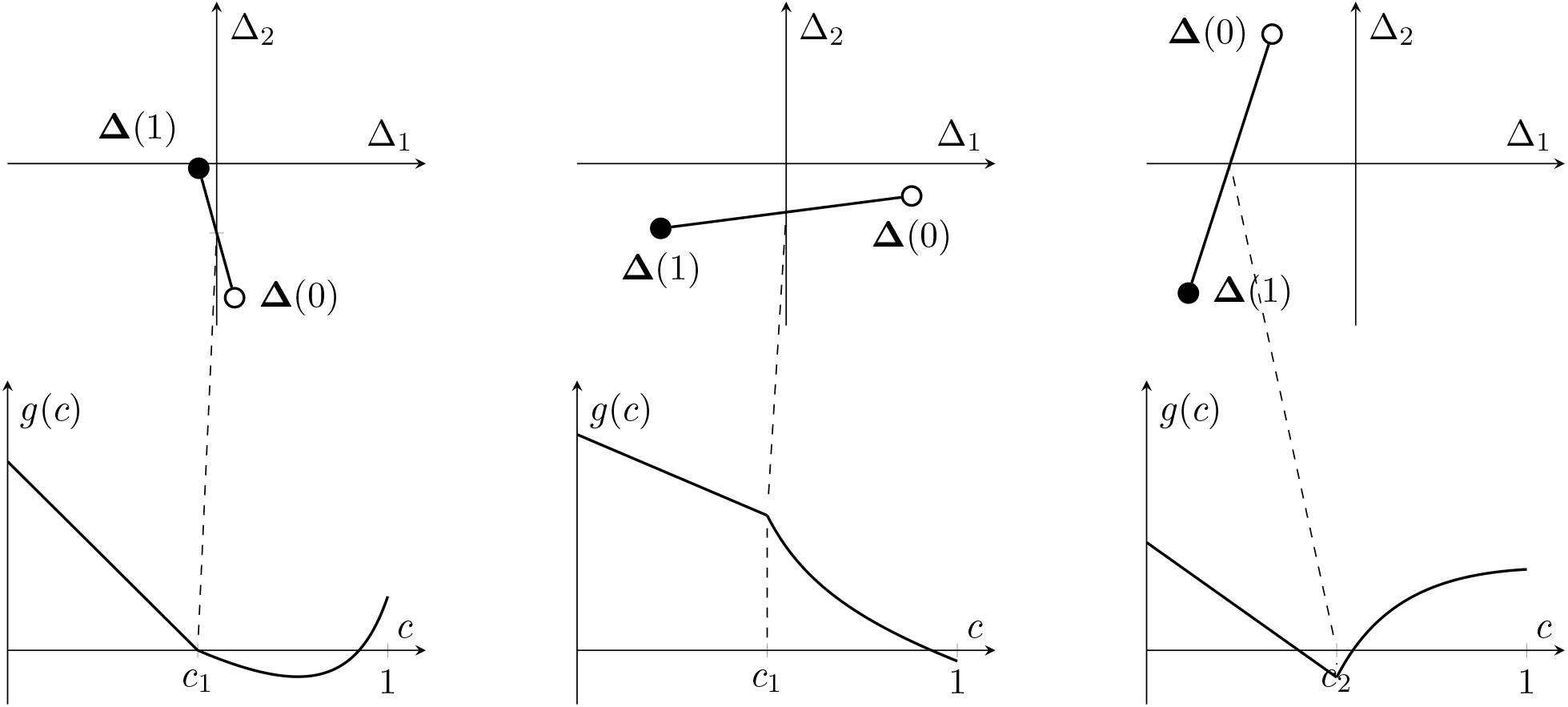
The dilemma strengths and equilibrium growth rates of the three games of Table 4. Equilibrium growth rate is a convex (left), concave-convex (middle), and convex-concave (right) function of dosage.

### 3.3 Discontinuity of response in transitioning out of Coordination games

Uniquely among the four game classes, coordination games have a multiplicity of equilibria and therefore the selection dynamics of the tumor are characterized by bi-stability. Predictions on the tumor’s behavior in case of transitioning into or out of the *C* quadrant are thus contingent on the tumor’s composition prior to transition.

In the case of transitioning into *C* from a neighboring quadrant, *PD* or *H*, we leverage the fact that both have a unique pure equilibrium. Under the assumption that the game spends enough time in the neighboring quadrant, by the time an increased dosage moves the game into the *C* quadrant, the tumor comprises mostly of one type of cell. Given this history, selection in the coordination game only consolidates the tumor’s composition towards the same pure equilibrium. Growth in this equilibrium is a linear function of dose, either *γ*_11_(*c*) or *γ*_22_(*c*), hence we conclude that transitioning into the *C* quadrant comes with a proportional response in dose.

In the case of transitioning out of the *C* quadrant into a neighboring quadrant, *PD* or *H*, there are several sub-cases. If the tumor’s history is such that transition does not change equilibrium, that is, if the tumor was type 1 in the *C* quadrant before a transition to *H* or type 2 before a transition to *PD*, response is linear as before. If the tumor changes equilibrium, response is discontinuous and changes slope upon transition.

For a detailed analysis, consider a mostly type 2 tumor in the *C* quadrant characterized by growth game **Π** and assume that **Π** − **K** lies in the *H* quadrant. Then, the point of transition is a dose of *c*_2_ and realized equilibrium growth is given by

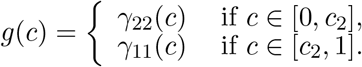

Equilibrium growth is linear in both the [0, *c*_2_] and [*c*_2_, 1] intervals, with a jump discontinuity in *c*_2_ from *γ*_22_(*c*_2_) to *γ*_11_(*c*_2_). At the same point the growth function changes slope from *κ*_22_ to *κ*_11_. We identify three cases:

1. if *γ*_22_(*c*_2_) *< γ*_11_(*c*_2_) and *γ*_22_(1) *< γ*_11_(1), equilibrium growth reduction is higher per dose unit for dose levels in [*c*_2_, 1] than in [0, *c*_2_].
2. if *γ*_22_(*c*_2_) *> γ*_11_(*c*_2_) and *γ*_22_(1) *> γ*_11_(1), equilibrium growth reduction is lower per dose unit for dose levels in [*c*_2_, 1] than in [0, *c*_2_].
3. otherwise equilibrium growth reduction may be higher or lower per dose unit in [*c*_2_, 1] than [0, *c*_2_] depending on the exact dosages.

In cases (1) and (2), the jump discontinuity and the changing of the slope of *g*(*c*) either act in the same direction or the latter effect is not large enough to offset the former. Thus, transitioning to the *H* quadrant increases per dose unit effectiveness in case (1) or lowers it in case (2). In case (3), the two act in opposition and the latter is large enough to offset the former for close to maximum doses. The effect of jump discontinuity dominate for doses close to *c*_2_, while for doses around 1, the difference in slope matters more.

### 3.4 Transitions through the coordination quadrant and response hysteresis loops

As shown in the previous section, transitioning into the *C* quadrant from one with a single equilibrium selects the dominant cell type of the tumor within the *C* quadrant. In addition, transitioning out of the *C* quadrant may switch equilibrium tumor composition. Both effects are present in case of a transition path between *PD* and *H* via *C* (Figure 6, top). In this case, the *C* quadrant can be entered both from the *PD* quadrant, in case of a dose increase, or the *H* quadrant, in case of a dose decrease. Thus, uniquely among the transition paths, a hysteresis loop in tumor growth rate is possible.

**Figure 5.**
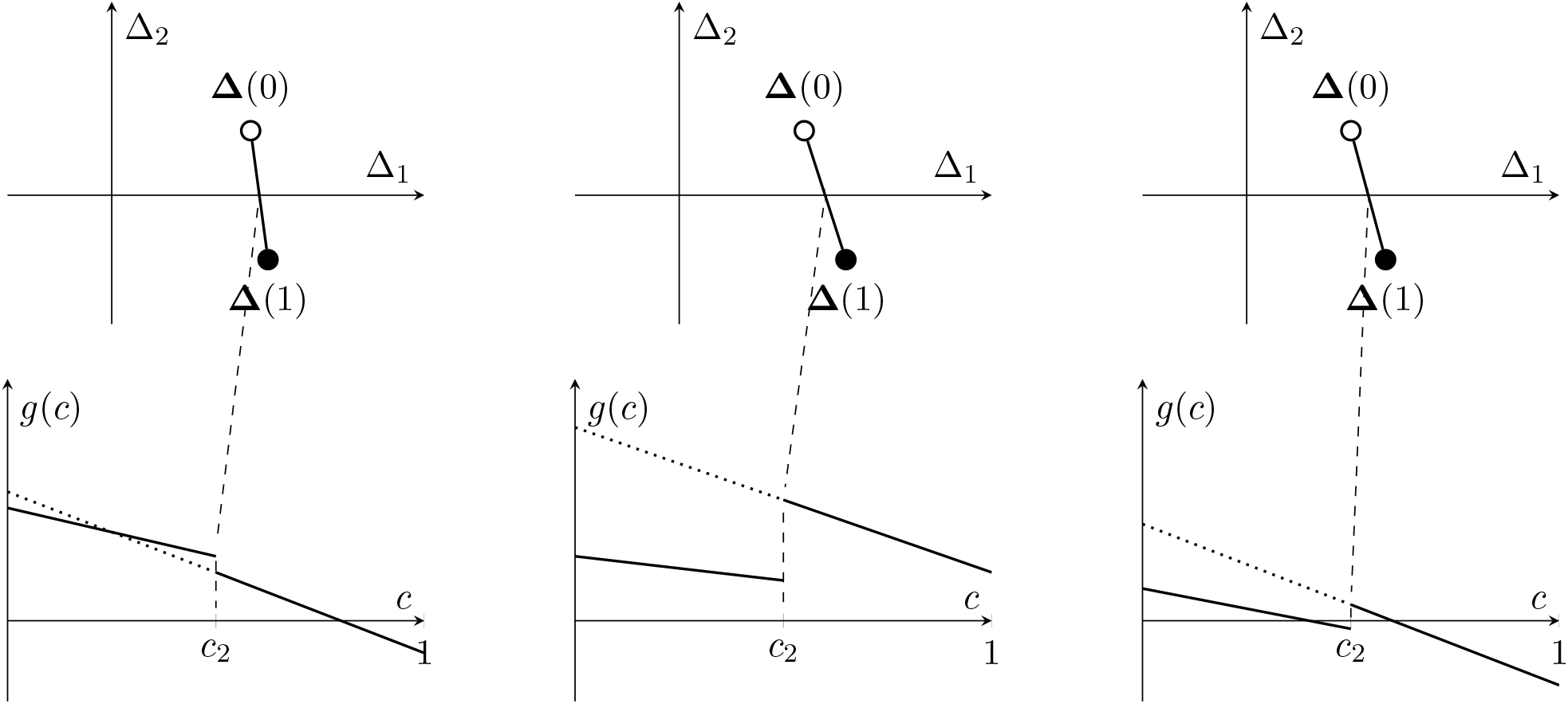
The dilemma strengths and equilibrium growth rates of the three games of Table 5 showing *C*-*H* transitions with *g*(*c*) shown in solid and the unrealized part of *G*(*c*) shown in dashed lines. After transition, response to treatment in terms of equilibrium growth reduction may be higher per dose unit may be higher (left), lower (middle), or may depend on the dose level post-transition (right): close to *c*_2_, response is lower, close to a maximum dose, it is higher.

**Figure 6.**
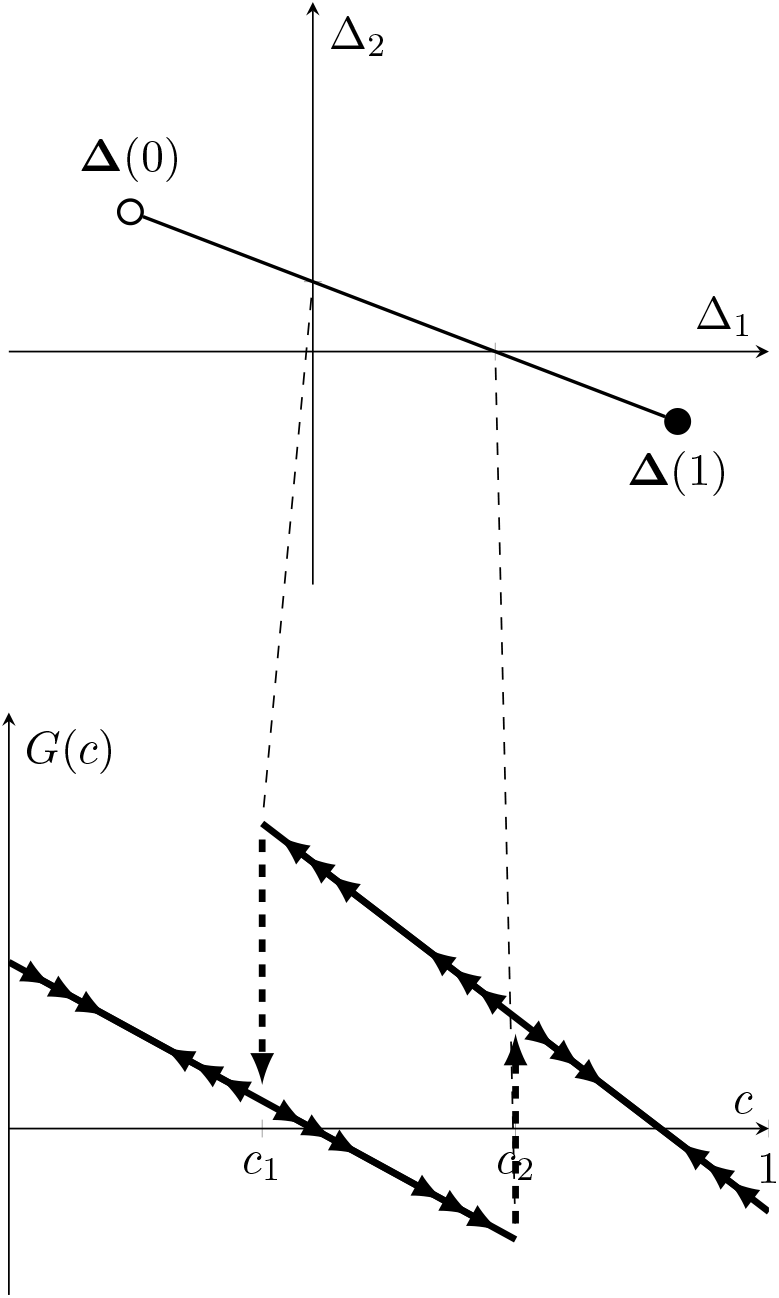
The hysterisis loop of equilibrium growth in a *PD*-*C*-*H* transition.

To illustrate, suppose that the game **Π** is in the *PD* quadrant and **Π** − **K** in the *H* quadrant such that *c*_1_ *< c*_2_. Under these conditions, there exists *c* ∈ [*c*_1_, *c*_2_] such that **Γ**(*c*) is in the *C* quadrant. Equilibrium growth as a set-valued function is given by

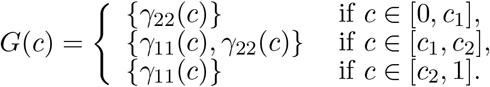

Consider a treatment strategy that ramps up dosage gradually from 0 to 1, then brings it back down to 0. The game moves from the *PD* quadrant to *H*, transiting through *C*, then back. Equilibrium growth along this trajectory will trace *γ*_22_(*c*) as dose is on the rise, until it reaches *c*_2_. At that point, equilibrium growth jumps to the *γ*_11_(*c*) line. Once dose is maximized and begins to be brought back down, equilibrium growth traces *γ*_11_(*c*) until dose reaches *c*_1_, then jumps to *γ*_22_(*c*). Equilibrium growth rate for dose levels *c* ∈ (*c*_1_, *c*_2_) is decided by which quadrant the game visited before entering *C*. A game showing a *PD*-*C*-*H* transition is shown in Table 6 while the hysteresis loop is showcased in Figure 6.

**Table 6.** Game and treatment parameters defining a transition from *PD* to *H* via *C*.

| $\Pi^{\text{PCH}}$ | 1 | 2 | $K^{\text{PCH}}$ | 1 | 2 |
| --- | --- | --- | --- | --- | --- |
| 1 | 8 | 6 | 1 | 10 | 3 |
| 2 | 0 | 7 | 2 | 11 | 5 |

## 4 Discussion

In this paper we model the selection and growth dynamics of cancer by a 2 × 2 evolutionary game. To obtain a classification of such games, we adopt the concept of ‘dilemma strength’ from the evolutionary game theory social dilemma literature, distinguishing four game classes by the signs of the differences of payoffs in the same column of the game matrix. The classes encode all information about the game’s equilibrium structure.

We then consider the effect of treatment. Treatment lowers the nominal payoff parameters of the game, inducing a first-order effect of reduced growth, as well as changing relative payoffs, inducing a second-order effect through altered selection dynamics. Despite a simple assumption of a linear dose effect in game-space (a reasonable first approximation upon surveying results of the evolutionary game assays in [30] and [31]), this mathematical framework allows for a wide range of outcomes in equilibrium growth-function space. Equilibrium growth dynamics are linear within the Prisoner’s Dilemma, Coordination, and Harmony classes and second-order effects are nil. In the Anti-Coordination quadrant, equilibrium growth may exhibit either convexity or concavity. As observed by measuring effective games in cancer, treatment may change the game class as well, further enriching the types of secondorder effects possible: transitioning into or out of the Anti-Coordination quadrant may produce convex, convex-concave, concave-convex, or concave equilibrium growth rates. Transitioning into the Coordination quadrant may change the type of equilibrium selected, creating a discontinuity and changed slope in equilibrium growth. Finally, our framework exhibits hysteresis, illustrating a path dependence in dosing with games passing through the Coordination quadrant.

The value of this exercise is two-fold. First, we hope to provide a unified and simple taxonomy for defining 2 × 2 effective games in cancer, replacing existing ones in use originally created for non-cooperative games. The two-dimensional representation of the four possible game-classes allows for a visual guide in mapping the relation and possible transitions between classes. For instance, the transition shown in [30] from Prisoner’s Dilemma to Harmony implies that lower treatment dose ought to land the game in one of the Coordination or Anti-Coordination quadrants, creating either bi-stability or coexistence of parental and resistant cell types. Second, we propose a relationship between the parameters characterizing cancer selection and growth and the types of second-order treatment effects. A number of discussion points are in order highlighting our model’s scope and limitations. First, in this study we make inferences on treatment convexity in terms of the game’s equilibrium growth rate. Under unconstrained growth, the equilibrium of the game is the asymptotic limit of the tumor’s composition, but whether the realities of the disease and the treatment make it a close approximation of realized cancer growth in vitro or in vivo is not straightforward. Without a change in game class, our simulations confirm that linearity of treatment response (e.g. prisoner’s dilemma game) leads to approximately equivalent outcomes, while a convex of response (e.g. an anti-coordination game) leads to the superiority of even treatment schedules (Figure 3).

Second, we restrict our analysis to frequency-dependent selection and growth excluding density or population effects such as carrying capacity or exogenous cell death. This assumption makes our model tractable, enabling us to study the effects of in changes in frequency-dependent selection of cancer in isolation. Increasing the richness of the model in this way would, among other effects, impact the linearity of transitions in dilemma-strength space. As the observations of [31] and [30] are consistent with linearity, restricting attention to frequency-dependence does not seem, at first glance, to be costly in terms of model accuracy.

Third, our attention is on a single therapy, defining one vector in dilemma-strength space. Allowing for multiple types of therapy would define the same number of vectors. In theory, any point in the span of vectors would be reachable by mixtures of therapies, expanding the set of transitions available for a given disease parametrization. Applying different therapies in succession creates paths in dilemma strength-space composed of concatenations of vectors, allowing for complicated sequences of transitions.

## Footnotes

1 Each of the four games correspond to an archetypal psychological pressure that the authors dubbed “exploiter” (a change of strategy affects self positively, has negative effects on other), “leader” (positive for both, higher for self), “hero” (positive for both, higher for other), and “martyr” (negative for self, positive for other, coinciding with the Prisoner’s Dilemma)

2 Strict means that indifferences are ruled out, ordinal means that games are considered equivalent up to monotone transformations of the payoffs.

3 We note that our Δ_1_ is the negative of the ‘gamble-intending-dilemma’ while Δ_2_ equals the ‘risk-averting-dilemma’.

4 The authors of [30] do not name the name classes, whereas [31] calls an anti-coordination game a ‘Leader game’, following [12]. We argue that this classification is not necessarily a good fit for evolutionary games. In behavioral applications of non-cooperative games, games may be considered equivalent up to a relabeling of one or both players’ strategies. Coordination games and anti-coordination games are equivalent up to a relabeling of only one player’s strategies; [12] calls their ‘leader game’ a variant of the ‘battle of the sexes’ game, an asymmetric coordination game. In evolutionary games with no relabeling allowed, ‘leader games’ and ‘hero games’ are always anti-coordination games, whereas symmetric coordination games are not covered in this classification, as [12] considers them to be trivial. Citing the nomenclature of [13], [31] correctly calls a harmony game ‘deadlock’, but this topological classification does not distinguish between ‘prisoner’s dilemma’ and ‘deadlock’.

5 Note that, because the labeling of the quadrants is made in terms of the growth game Π, interaction games in *H* or *PD* quadrant may not be harmony or prisoner’s dilemma games. For example, a game Γ(*c*) where type 1 is dominated is in the *PD* quadrant as type 1 is growth-optimal without treatment, but *γ*_11_(*c*) *< γ*_22_(*c*) may obtain, in which case Γ(*c*) is not itself be a true prisoner’s dilemma game. Similarly, a game Γ(*c*) in the *H* quadrant is not a true harmony game itself unless *γ*_11_(*c*) *> γ*_22_(*c*) holds. Interaction games in the *AC* or the *C* quadrant, however, are always indeed anti-coordination and coordination games, respectively.

6 Note: linearity (or convexity/concavity) serves as only an approximation of outcomes due to the fact that tumors may be initially far from equilibrium growth dynamics (see Discussion for more).

